# Synthetic Heparan Sulfate Mimetic Pixatimod (PG545) Potently Inhibits SARS-CoV-2 By Disrupting The Spike-ACE2 interaction

**DOI:** 10.1101/2020.06.24.169334

**Authors:** Scott E. Guimond, Courtney J. Mycroft-West, Neha S. Gandhi, Julia A. Tree, Thuy T Le, C. Mirella Spalluto, Maria V. Humbert, Karen R. Buttigieg, Naomi Coombes, Michael J. Elmore, Kristina Nyström, Joanna Said, Yin Xiang Setoh, Alberto A. Amarilla, Naphak Modhiran, Julian D.J. Sng, Mohit Chhabra, Paul R. Young, Marcelo A. Lima, Edwin A.Yates, Richard Karlsson, Rebecca L. Miller, Yen-Hsi Chen, Ieva Bagdonaite, Zhang Yang, James Stewart, Edward Hammond, Keith Dredge, Tom M.A. Wilkinson, Daniel Watterson, Alexander A. Khromykh, Andreas Suhrbier, Miles W. Carroll, Edward Trybala, Tomas Bergström, Vito Ferro, Mark A. Skidmore, Jeremy E. Turnbull

**Affiliations:** Molecular & Structural Biosciences, School of Life Sciences, Keele University, Newcastle-Under-Lyme, Staffordshire, ST5 5BG, UK; School of Chemistry and Physics, Centre for Genomics and Personalized Health, Queensland University of Technology, 2 George Street, Brisbane, QLD 4000, Australia; National Infection Service, Public Health England, Porton Down, Salisbury, Wiltshire, England, UK, SP5 3NU; Queensland Institute of Medical Research Berghofer Medical Research Institute, Brisbane, Queensland 4029, Australia; School of Clinical and Experimental Sciences, University of Southampton Faculty of Medicine, Southampton, UK; Department of Infectious Diseases, Institute of Biomedicine, University of Gothenburg, Guldhedsgatan 10B, S-413 46 Goteborg, Sweden; School of Chemistry and Molecular Biosciences, University of Queensland, Brisbane, QLD 4072, Australia; Australian Infectious Diseases Research Centre, University of Queensland, Brisbane, QLD 4072, Australia; Department of Biochemistry and Systems Biology, Institute of Systems, Molecular and Integrative Biology, University of Liverpool, Liverpool, L69 7ZB, UK; Copenhagen Center for Glycomics, Department of Cellular & Molecular Medicine, University of Copenhagen, Copenhagen N 2200, Denmark; Dept of Infection Biology & Microbiomes, University of Liverpool, Liverpool, L69 7ZB, UK; Zucero Therapeutics Ltd, 1 Westlink Court, Brisbane, Queensland 4076, Australia; NIHR Southampton Biomedical Research Centre, University Hospital Southampton, UK

## Abstract

Heparan sulfate (HS) is a cell surface polysaccharide recently identified as a co-receptor with the ACE2 protein for recognition of the S1 spike protein on SARS-CoV-2 virus, providing a tractable new target for therapeutic intervention. Clinically-used heparins demonstrate inhibitory activity, but world supplies are limited, necessitating alternative solutions. Synthetic HS mimetic pixatimod is a drug candidate for cancer with immunomodulatory and heparanase-inhibiting properties. Here we show that pixatimod binds to and destabilizes the SARS-CoV-2 spike protein receptor binding domain (S1-RBD), and directly inhibits its binding to human ACE2, consistent with molecular modelling identification of multiple molecular contacts and overlapping pixatimod and ACE2 binding sites. Assays with multiple clinical isolates of live SARS-CoV-2 virus show that pixatimod potently inhibits infection of monkey Vero E6 and human bronchial epithelial cells at concentrations within its safe therapeutic dose range. Furthermore, in a K18-hACE2 mouse model pixatimod demonstrates that pixatimod markedly attenuates SARS-CoV-2 viral titer and COVID-19-like symptoms. This demonstration of potent anti-SARS-CoV-2 activity establishes proof-of-concept for targeting the HS-Spike protein-ACE2 axis with synthetic HS mimetics. Together with other known activities of pixatimod our data provides a strong rationale for its clinical investigation as a potential multimodal therapeutic to address the COVID-19 pandemic.

## Introduction

The coronavirus disease 2019 (COVID-19) pandemic caused by the severe acute respiratory syndrome coronavirus 2 (SARS-CoV-2) has according to the World Health Organisation recently surpassed 110 million cases and 2.4 million deaths world-wide. Although vaccines against COVID-19 are currently being developed and deployed, given the severe pathophysiology induced by SARS-CoV-2 (*1*), there is a clear need for therapeutic alternatives to alleviate and stop the COVID-19 epidemic that complement vaccination campaigns. Heparan sulfate (HS) is a highly sulfated glycosaminoglycan found on the surface of most mammalian cells which is used by many viruses as an entry receptor or co-receptor (*2*), including coronaviruses (*3*). Various compounds that mimic cellular HS such as clinically-used heparins have been investigated and have been shown to block infectivity and cell-to-cell spread in a multitude of different viruses, including SARS-associated coronavirus strain HSR1 (*4*). The glycosylated spike (S) protein of SARS-CoV-2 mediates host cell infection via binding to a receptor protein, angiotensin-converting enzyme 2 (ACE2) (*5*). Analysis of the sequence and experimentally determined structures of the S protein reveals that the receptor binding domain (RBD) of the S1 subunit contains a HS binding site. Recent studies have clearly demonstrated binding of heparin and HS to S1 RBD (*6-9*), including induction of significant conformational change in the S1 RBD structure (*6*), and also revealed that HS is a co-receptor with ACE2 for SARS-CoV-2 (*10*). Collectively these data strongly suggest that blocking these interactions with heparins and HS mimetics has potential as an effective strategy for COVID-19 therapy. Although heparins have major potential for repurposing for such applications, limitations in the global supply of natural product heparins will greatly restrict its availability (*11*), highlighting an urgent need to find synthetic alternatives.

Pixatimod (PG545) is a clinical-stage HS mimetic with potent anti-cancer (*12,13*), and anti-inflammatory properties (*14*). However, significant antiviral and virucidal activity for pixatimod has also been reported against a number of viruses that use HS as an entry receptor with EC_50_’s ranging from 0.06 to 14 µg/mL. This includes HSV-2 (*15*), HIV (*16*), RSV (*17*), Ross River, Barmah Forest, Asian CHIK and chikungunya viruses (*18*), and Dengue virus (*19*). *In vivo* efficacy has been confirmed in a prophylactic mouse HSV-2 genital infection model (*15*), a prophylactic Ross River virus mouse model (*18*) and a therapeutic Dengue virus mouse model (*19*). Pixatimod has been evaluated in a Phase Ia clinical trial in patients with advanced solid tumours where it demonstrated a tolerable safety profile and some evidence of disease control (*13*). It has been safely administered to over 80 cancer patients in Phase I studies as a monotherapy or in combination with nivolumab (NCT02042781 and ACTRN12617001573347), prompting us to examine its anti-viral activity against SARS-CoV-2.

Here we provide evidence of a direct and destabilizing interaction of pixatimod with the S1 spike protein RBD, supported by molecular modelling data. Additionally, pixatimod was able to inhibit the interaction of S1-RBD with ACE2 and also Vero cells which are known to express the ACE2 receptor, indicating a direct mechanism of action. We established that pixatimod is a potent inhibitor of attachment and invasion of Vero cells and human bronchial epithelial cells by multiple clinical isolates of live SARS-CoV-2 virus, and reduces its cytopathic effect, at concentrations within the known therapeutic range of this drug. Finally, we observed marked attenuation of SARS-CoV-2 viral RNA load and COVID-19-like symptoms in the K18-hACE2 mouse model of infection. Our data demonstrate that synthetic HS mimetics can target the HS-Spike protein-ACE2 axis to inhibit SARS-CoV-2 infection. They provide strong support for clinical investigation of the potential of pixatimod as a novel therapeutic intervention for prophylaxis and treatment of COVID-19, and have implications for wider applications against other HS-binding viruses and emerging global viral threats.

## Results

### Modelling of pixatimod-spike protein interactions

We initially used molecular dynamics (MD) simulations to map the potential binding sites of pixatimod (**Fig 1A**) on the S1 RBD surface (**Fig 1B**) of monomeric spike. A total of 24 unique residues of RBD are known to interact with ACE2 based on the X-ray structures (**Fig 1B**). Interestingly, a few of these residues (Tyr489, Phe456, Leu455, Ala475) are also predicted to be involved in binding to pixatimod. Amino acids making significant interactions with pixatimod were identified on the basis of their individual contributions to the total interaction energy, considering only the residues that contribute less than -1.0 kcal/mol. The decomposition approach was helpful for locating residues of the RBD domain such as Lys458, Ser459, Lys462 and Asn481 that transiently interact to form hydrogen bonds or ionic interactions with the sulfated tetrasaccharide moiety of pixatimod (**Fig 1D**). The free energy of binding is -10 kcal/mol, wherein van der Waals energies make the major favourable contribution to the total free energy. The cholestanol residue also formed stabilizing interactions with Tyr489, Phe456, Tyr473, Ala475, Gln474 and Leu455 (**Fig 1E**). Furthermore, the standard deviation of backbone RMSD around residues Leu455-Pro491 and the N-terminal of RBD (Thr333-Thr345) among the four repeated MD trajectories were approximately 2Å, indicating significant conformational change in the region. RMSF calculations of main-chain atoms showed significant atomic fluctuations (≥1.5 Å) for Lys458, Asn460, Lys462, Arg466, Ser477 and Asn481 upon binding to the ligand pixatimod. These results indicate that a conformational change may be induced by binding of pixatimod to S1 RBD (**Fig 1C**).

**Figure 1:**
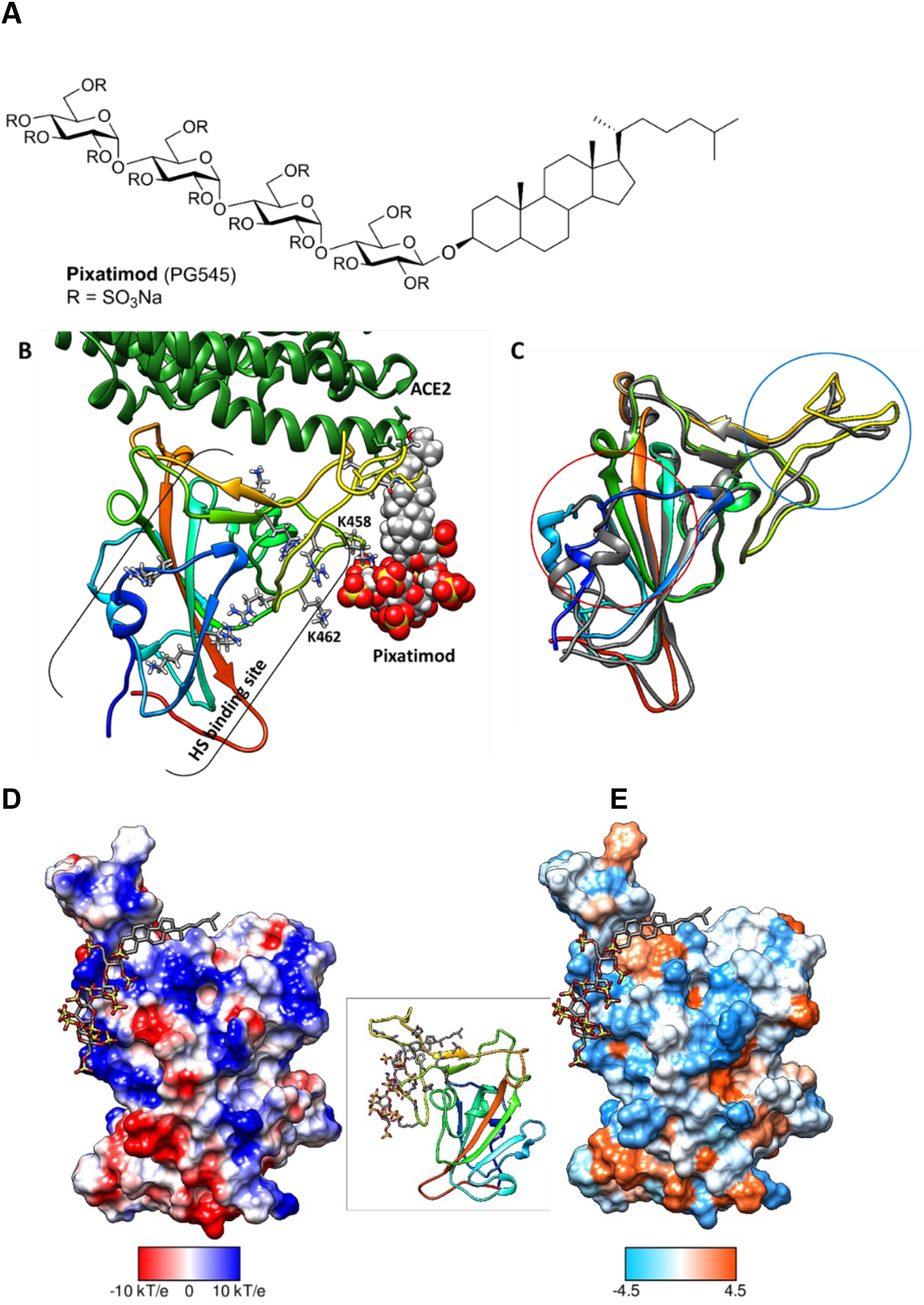
Molecular dynamics modelling defines direct interactions of pixatimod with S1 RBD: **A**, Structure of pixatimod. **B**, Model showing interactions of pixatimod with the RBD domain of spike protein. The sulfated tetrasaccharide partially occupies the HS/heparin binding site. The lipophilic tail of pixatimod wraps around the hydrophobic loop, thereby creating a steric clash with the helix of ACE2 protein (shown in inset-green ribbon). **C**, Superimposition of the X-ray structure (PDB: 6LZG) and one of the snapshots from MD simulations (ligand not shown) suggest conformational change around the loop region (blue circle) and the N-terminal helix as highlighted (red circle). **D**, coulombic surface and **E**, hydrophobic surface binding mode of pixatimod to S1 RBD. Both surfaces are oriented in the same direction as shown in the ribbon diagram of the protein in the middle. The sulfated tetrasaccharide interacts with the basic regions on S1 RBD whereas cholestanol residue prefers hydrophobic region for interactions. Coulombic surface coloring defaults: ε = 4r, thresholds ± 10 kcal/mol were used. Blue indicates surface with basic region whereas red indicates negatively charged surface. The hydrophobic surface was colored using the Kyte-Doolittle scale wherein blue, white and orange red colour indicates most hydrophilicity, neutral and hydrophobic region, respectively. UCSF Chimera was used for creating surfaces and rendering the images. Hydrogens are not shown for clarity.

An alternate heparin binding site is reported around residues Arg403, Arg406, Arg408, Gln409, Lys417, Gln493, Gln498 (*8*). One of the replicates indicated a second binding mode wherein the tetrasaccharide of pixatimod was found to interact around this region (Supplementary Materials, **Fig S1**), however, the free energy of binding was > +13 kcal/mol indicating much less favourable binding to this site. Overall, our modelling data strongly support the notion of direct binding of pixatimod via multiple amino acid contacts in S1 RBD, potentially resulting in induction of a conformational change and interference with binding to ACE2.

### Pixatimod interacts with spike protein

Spectroscopic studies with circular dichroism (CD) were undertaken to investigate direct binding of pixatimod to recombinant spike protein receptor binding domain (S1 RBD) (**Fig 2A,B**), the region which interacts with the ACE2 receptor on human cells. CD spectroscopy in the far UV region (λ = 190 - 260 nm) detects conformational changes in protein secondary structure that occur in solution and can infer binding by an added ligand. Such secondary structural changes can be quantified using spectral deconvolution. SARS-CoV-2 EcS1-RBD (produced in *E. coli*, see Methods) underwent conformational change in the presence of either pixatimod or heparin as a comparator sulfated molecule known to bind the RBD (*6-10*), consisting of decreased α-helical content for pixatimod and increased α-helical content for heparin (**Fig 2C**). A decrease in global β-sheet content is observed for both pixatimod and heparin, along with increases in turn structure (**Fig 2C**).

**Figure 2:**
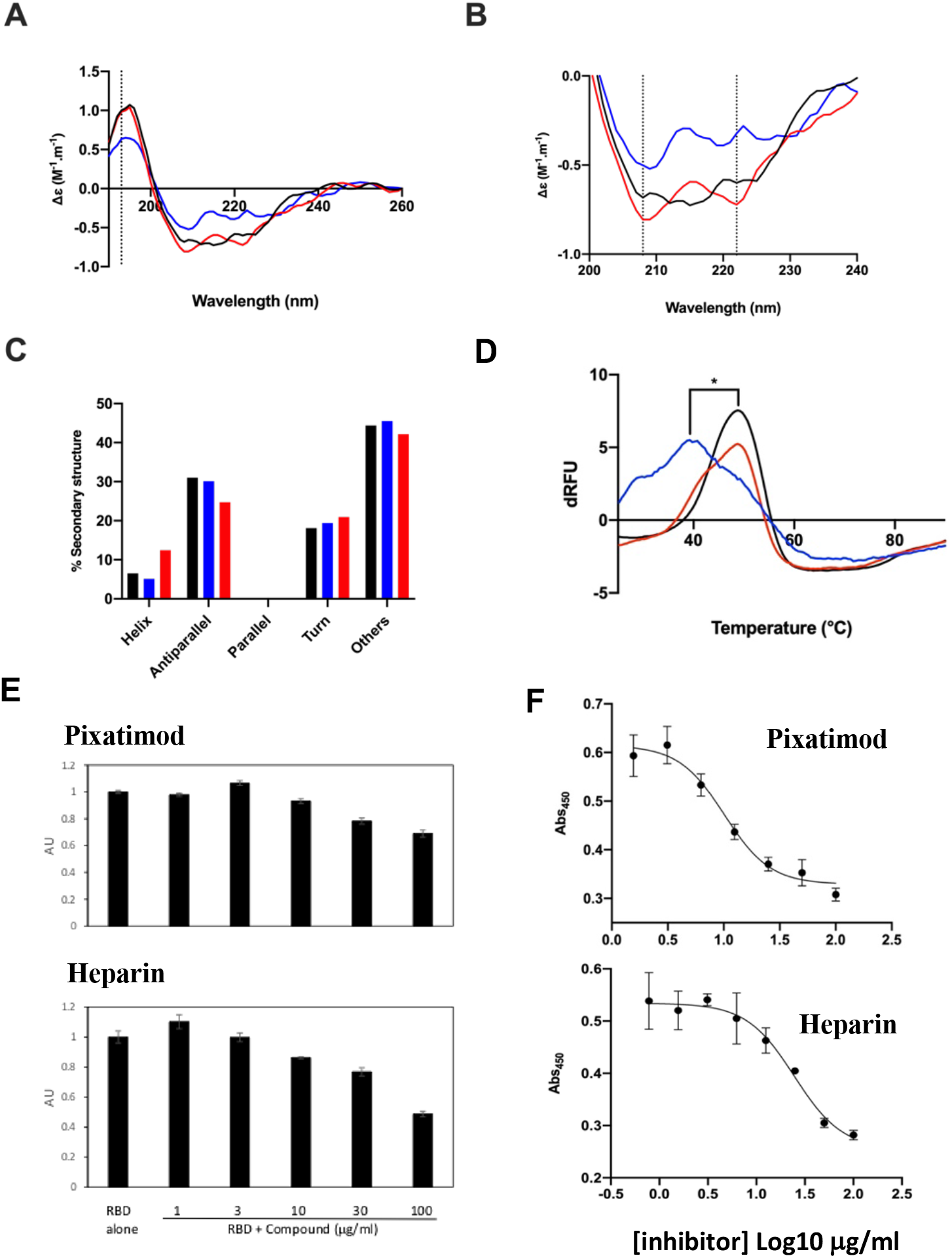
Pixatimod interacts with SARS-CoV-2 S1-RBD and inhibits binding to cells and ACE2 receptor. **A**, Circular dichroism spectra (190 - 260 nm) of SARS-CoV-2 EcS1-RBD alone (black), or with pixatimod (blue) or heparin (red). The vertical dotted line indicates 193 nm. **B**, The same spectra expanded between 200 and 240 nm. Vertical dotted lines indicate 222 nm and 208 nm. **C**, Secondary structure content analysed using BeStSel for SARS-CoV-2 EcS1-RBD. α-helical secondary structure is characterized by a positive band at ∼193 nm and two negative bands at ∼208 and ∼222 nm (analysis using BeStSel was performed on smoothed data between 190 and 260 nm). **D**, Differential scanning fluorimetry of binding of pixatimod (blue; 10μg) or heparin (red; 10 μg) to mS1-RBD (1μg; black line, protein-only control). *T_m_ values for RBD alone (48.4°C, SD = 0.3) and in the presence of pixatimod (39.3°C, SD = 1) were statistically different, *t*(4) = 15.25, *p* = 0.0001. **E**: dose response effects of pixatimod (E) and unfractionated porcine mucosal heparin (F) on binding of EcS1-RBD to Vero cells. Data were normalised to control with no addition of EcS1-RBD. AU, arbitrary units of fluorescence. n=3 +/- CV. **F**, Competitive ELISA assay using biotinylated human ACE2 protein immobilized on streptavidin coated plates, to measure inhibition of binding of mS1-RBD in the presence of various concentrations of inhibitor compounds. Pixatimod (IC_50_, 10.1 μg/ml) and porcine mucosal heparin (IC_50_, 24.6 μg/ml). n=3, +/-SD; representative example shown.

We explored the effects of pixatimod on protein stability using differential scanning fluorimetry (DSF) in which the thermal denaturation of a protein is monitored in the presence of a hydrophobic fluorescent dye (*20*). Binding of pixatimod induced a notably large reduction in melting temperature (ΔT_m_) of 9.1°C (**Fig 2D**; p=0.0001), indicating major destabilisation of the mammalian expressed S1-RBD (mS1 RBD) protein. In contrast, heparin at an equivalent dose only partially destabilised the RBD protein, evidenced by a small side peak shifted by ∼5-6°C indicating populations of RBD in a bound and unbound state (**Fig 2D**).

The observed changes demonstrate that the SARS-CoV-2 S1 RBD interacts with pixatimod in aqueous conditions of physiological relevance. Notably, the conformational changes and destabilization observed were distinct for pixatimod compared to heparin, suggesting distinct interactions (**Fig 2**). Consistent with the modelling results, these data confirm direct interactions of pixatimod with S1 RBD, resulting in induction of a conformational change, consistent with the notion that HS mimetics such as pixatimod have the potential to interfere with S1-RBD interactions with ACE2.

### Pixatimod inhibits S1-RBD cell-binding

We next evaluated inhibition of binding of His-tagged EcS1-RBD to monkey Vero cells (which are known to express both HS proteoglycans (HSPGs) and the ACE2 protein receptor required for SARS-CoV-2 attachment and cell invasion). Fixed cells were exposed to His-tagged S1 RBD for 1hr, in the presence or absence of additional compounds, with subsequent washing and detection using a fluorescently-labelled anti-His tag antibody. A clear dose response was noted for both pixatimod and heparin as a comparator compound (**Fig 2E**), with 32% and 51% inhibition achieved at 100 µg/mL respectively. These data confirm that pixatimod can interfere with binding of S1-RBD to cells containing HSPGs and ACE2 protein receptors.

### Pixatimod directly inhibits S1-RBD binding to ACE2

To further evaluate the mechanism of action of pixatimod its direct effects on the interaction of S1-RBD with the ACE2 protein receptor was measured using a competitive ELISA assay. Inhibition of binding of mS1-RBD preincubated with various concentrations of inhibitor compounds was measured by detection with an anti-RBD antibody. A dose response was observed with pixatimod showing an IC_50_ of 10.1 μg/ml (**Fig 2F**). In comparison heparin also demonstrated inhibitory activity but with lower potency (24.6 μg/ml; **Fig 2F**). Importantly, this data confirms a direct mechanism of action of pixatimod via inhibition of S1-RBD binding to the ACE2 protein receptor.

### Pixatimod inhibits SARS-CoV-2 infection

The effect of pixatimod on SARS-CoV-2 infection of Vero cells was examined using a standard plaque reduction neutralisation assay. Pixatimod was pre-incubated with the SARS-CoV-2 clinical isolate from Victoria, Australia (VIC01) for 1 hr before infecting the cells. Significant decreases were observed in the number of PFU after pixatimod treatment of SARS-CoV-2 **(Fig 3A**). Analysis of multiple dose response curves yielded an EC_50_ for pixatimod in the range of 2.4-13.8 μg/mL (mean 8.1 μg/ml; n=3 assays) (**Table 1**). In comparison, an EC_50_ of ∼10 μg/ml has been observed for unfractionated heparin with a SARS-CoV-2 Italy UniSR1/2020 isolate (*8*) and 20-64 μg/ml for the SARS-CoV-2 VIC01 isolate (*21*).

**Figure 3:**
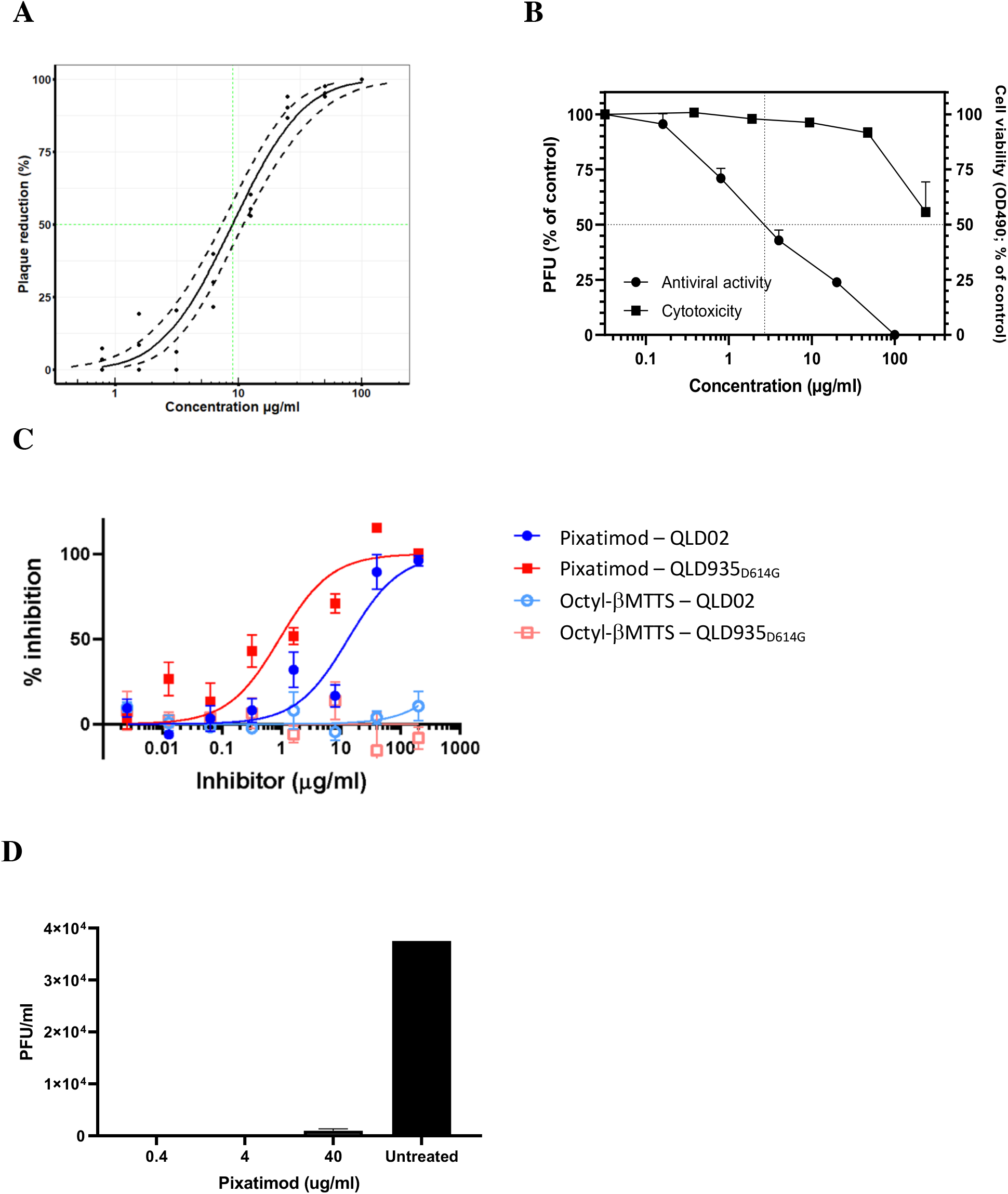
Pixatimod inhibits infection of Vero and human bronchial epithelial cells with different SARS-CoV-2 virus isolates. Live virus infectivity assays were performed as described in Methods for 3 different SARS-CoV-2 isolates (representative data shown). **A**, Plaque reduction neutralization assay of Victoria isolate (VIC01) with a Probit mid-point analysis curve ±95% confidence intervals (dashed lines) (EC_50_ 8.9 μg/ml; 95% CI, 7-11; n=3). **B**, Plaque reduction assay of DE isolate, EC_50_ 2.7 µg/ml; n=3, +/- SD. **C**, Cytopathic assay of Queensland isolates, EC_50_ 13.2 (QLD02) and 0.9 (QLD935 with D614G mutation) µg/ml n=6, +/- SEM. Representative examples are shown in each case. Results of pixatimod inhibition of SARS-CoV-2 infectivity are expressed as percent plaque reduction (A), plaque number as a percent of control (B), or percent inhibition from cytopathic effect (C). Panel B also shows cytotoxicity data for Vero cells for calculation of CC_50_ value (>236 µg/ml). In panel C, data is also shown for octyl β-maltotetraoside tridecasulfate (Octyl-βMTTS; Supplementary Materials Fig S2), an analogue of pixatimod which lacks the steroid side-chain. **D**. Plaque assay to measure inhibitory effect of pixatimod on viral shedding in BCi-NS1.1 bronchial epithelial cells grown in an air liquid interface (ALI). ALI cultures were infected for 2h with SARS-CoV-2 (VIC01, MOI=0.2) previously preincubated for 1 hr at 37°C with 0.4, 4 and 40 μg/ml of pixatimod or HBSS for untreated. After 72 hours an apical rinse was performed with 200 μl of HBSS and 100 μl of the wash were used in a plaque assay in Vero E6 cells. Values are expressed as plaque forming unit (PFU)/ml, n= 2 +/- SD.

**Table 1:** Anti-SARS-CoV-2 activities of pixatimod in Vero cells.

| SARS-CoV-2 isolate | Assay Method | EC<sub>50</sub> (µg/mL)<sup>a</sup> | CC<sub>50</sub> (µg/mL) | Selectivity Index<sup>a</sup> |
| --- | --- | --- | --- | --- |
| VIC01 isolate | Plaque reduction | 8.1 ± 3.1 (2.4-13.8)<sup>a</sup> | >236<sup>d</sup> | 29 (>17 to >98<sup>a</sup>) |
| DE-Gbg20 isolate | Plaque reduction | 2.7<sup>b</sup> |  | >87 |
|  | Cytopathic effect | (0.8 – 11.6)<sup>c</sup> |  | >20 to >295 |
| QLD02 isolate | Cytopathic effect | 13.2 (8.0 – 21.6) |  | >17 |
| QLD935 isolate | Cytopathic effect | 0.9 (0.4 – 1.9) |  | >200 |

<sup>a</sup> Mean values and individual assay result ranges, and resulting selectivity index ranges, in brackets. <sup>b</sup> Mean EC<sub>50</sub> ±SE based on the data from three independent virus plaque reduction assays (PRNT<sub>50</sub> values). <sup>c</sup> EC<sub>50</sub> computed by the Reed and Muench formula based on the cytopathic effect assay. Range indicates upper (complete protection of cells) and lower (partial protection) limits of EC<sub>50</sub> estimation. <sup>d</sup> Cytotoxicity in Vero cells (determined at University of Gothenburg).

To establish that these antiviral effects were relevant for wider clinical viral isolates, assays were conducted with the isolate DE-Gbg20 from Sweden in a plaque reduction assay. Pixatimod inhibited infectivity of the DE-Gbg20 isolate with an EC_50_ value of 2.7 µg/mL (**Fig 3B**), similar to that found in experiments with the VIC01 isolate. Analysis of pixatimod cytotoxicity for Vero cells using a tetrazolium-based assay revealed that pixatimod decreased by 50% (CC_50_) the viability of Vero cells at concentration >236 µg/mL, i.e., well above the EC_50_ values observed in the plaque reduction assay (**Fig. 3B; Table 1**). Selectivity index (SI) values for pixatimod ranged from >17 to >98 for these assays.

In addition to the plaque reduction assays pixatimod inhibition of SARS-CoV-2 infectivity was assessed using assays that measured the cytopathic effects of the virus as an endpoint. Using the Swedish DE-Gbg20 isolate, and two Australian isolates from Queensland (QLD02 and QLD935), the EC_50_ values for pixatimod inhibition of SARS-CoV-2 infectivity were determined to be 0.8-11.6, 13.2 and 0.9 μg/mL, respectively (**Table 1**), values comparable with those observed for the plaque reduction assays (**Table 1**). We also noted that a pixatimod analogue octyl β-maltotetraoside tridecasulfate (without the steroid side-chain) (**Fig S2**) lacked efficacy for both QLD02 and QLD935 isolates (**Fig 3C**), demonstrating the importance of the steroid side-chain for activity, and supporting the notion of the sterol moiety promoting RBD switching position and enhancing availability of the heparin binding site. Notably, both DE-Gbg20 and QLD935 isolates contain the D614G mutation of the spike protein commonly present in recent isolates (**Table S1**) (*22*). The QLD935 isolate exhibited lower cytopathicity, which could partially contribute to the observed lower EC_50_ for pixatimod against this isolate.

To provide evidence of antiviral properties of pixatimod in a more physiologically relevant model for infection of human cells, we used a bronchial airway epithelial *in vitro* system of SARS-CoV-2 infection and replication. The hTERT transformed bronchial epithelial cell line BCi-NS1.1 differentiates into airway multi-ciliated cells when grown in Transwells at the air-liquid interface (ALI) cultures which show robust infection with SARS-CoV-2 when harvested 72 h after inoculation (*23,24*). To validate this model we firstly infected ALI bronchial cultures by adding SARS-CoV-2 (VIC01, MOI=0.2) on the apical side of the culture system and measured up to 1000 fold increase in viral RNA between 2h and 72h post-infection, measured by RT-PCR of viral RNA using primers for N2/N3 regions of the N gene of SARS-CoV-2 (**Fig S3, panel A**). To determine if pixatimod has inhibitory effects on SARS-CoV-2 infection and replication in this model, the virus was preincubated with 0.4, 4 or 40 μg/ml of pixatimod (or HBSS for untreated) for 1 hr at 37°C before the inoculum was added to the apical side of the cells. We examined infectious virus in the apical wash from BCI ALI cultures by plaque assay on Vero E6 cells. Viral shedding of ALI cultures infected with SARS-CoV-2 in the presence of 0.4 and 4 μg/ml pixatimod was completely abolished in comparison to the cells infected with the untreated virus as a control (**Fig 3D**), whereas at 40 μg/ml pixatimod resulted in >40 fold reduction in PFU/ml **(Fig 3D)**. To corroborate these findings viral shedding from the apical side of the culture and viral load in the infected cells was also measured by RT-PCR. Incubation of SARS-CoV-2 with different concentrations of pixatimod reduced SARS-CoV-2 shedding from ALI cells into the apical phase by > 98%, and reduced SARS-CoV-2 RNA within the cells by >97% at 72hr at all pixatimod doses, compared to untreated infected cells **(Fig S3, panels A & B)**. Notably, efficacy at 0.4 and 4 μg/ml was >99% in both cases indicating high potency of pixatimod in inhibition of SARS-CoV-2 infection of human bronchial cells. Overall, these data in a physiologically-relevant human bronchial cell model show that pixatimod is a potent inhibitor of SARS-CoV-2 in terms of both lowering viral shedding in the apical compartment and viral load in the cell layer.

### Pixatimod inhibits SARS-CoV-2 infection *in vivo*

The potential efficacy of pixatimod *in vivo* was explored with an established SARS-CoV-2 lethal infection model using hACE2 transgenic mice wherein hACE2 is expressed behind a keratin 18 promoter (*25-28*). Virus was inoculated in 50 µl via the intranasal route into the lungs (*25-27*). A single prophylactic treatment of 16 mg/kg of pixatimod was given to K18-hACE2 mice one day before intranasal infection with SARS-CoV-2. Two control groups were included, K18-hACE2 mice that were treated with saline and infected with virus, and C57BL/6J mice given pixatimod but no virus. Mice were weighed daily and euthanized on day 5 when weight loss approached 20%, the ethically defined endpoint. As reported previously in other preclinical models of infectious disease (*18,19*), pixatimod led to an initial bodyweight loss of 8%. Weight loss was thus presented relative to day 1 post-infection in order to focus on infection-induced weight loss (**Fig. 4A**), which generally occurs after day 3 (*26*). The mean weight loss for saline-treated K18-hACE2 mice was significantly higher than for pixatimod treated K18-ACE2 mice (p=0.043, repeat measures ANOVA including data from days 4 and 5), with the latter showing no significant infection-associated weight loss (**Fig. 4A**).

**Fig 4:**
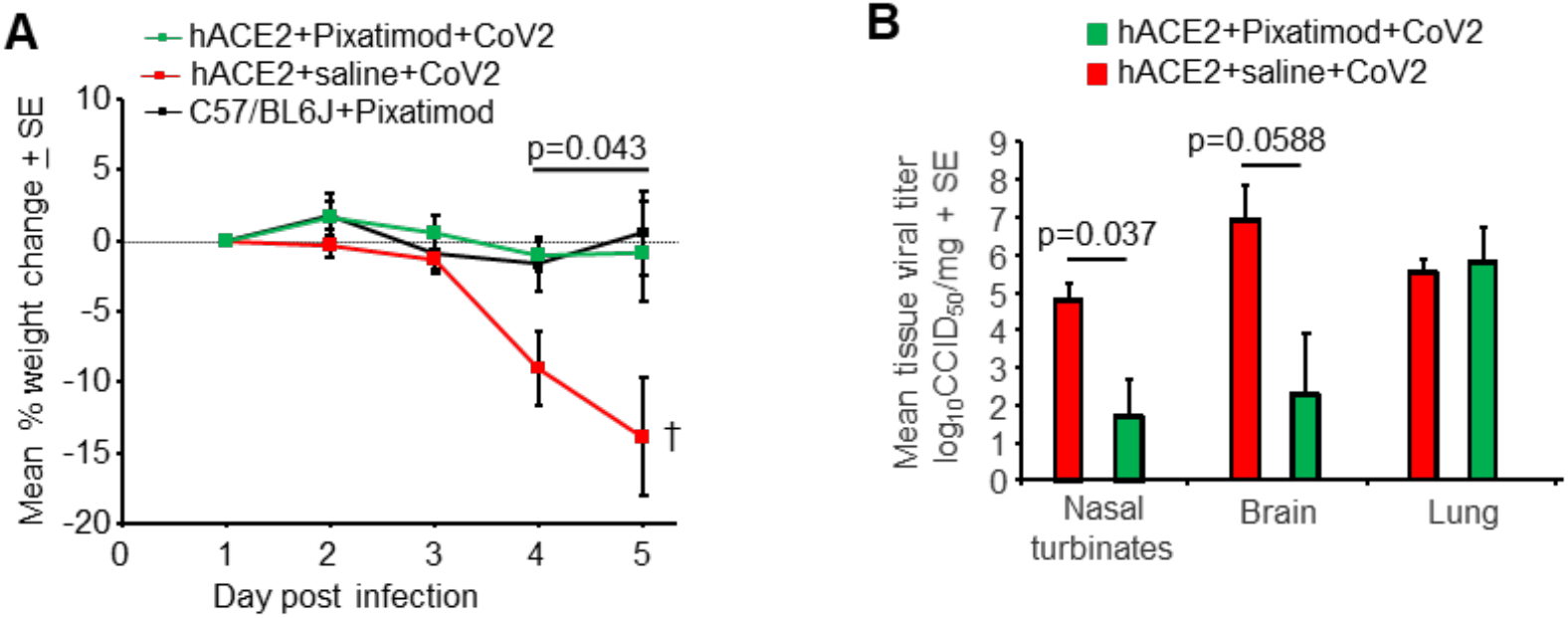
Pixatimod inhibits SARS-CoV-2 infection in K18 hACE2 transgenic mice. (A) Mean percentage weight change relative to day 1 post infection. Statistics by repeat measures ANOVA for days 4 and 5. n=4 mice per group, +/- SE. Mice were euthanized on day 5. (B) Mean tissue titres on day 5 post infection. Statistics by Kolmogorov Smirnov test (Nasal turbinates) and t test (Brain). n=4 mice per group, +/- SE.

Tissue viral loads were assessed on day 5 post infection, with pixatimod-treated animals showing a significant 3.1 log_10_ fold reduction in viral titers in nasal turbinates (p=0.037, Kolmogorov Smirnov test, non-parametric data distribution), and a 4.6 log_10_ fold reduction in viral titers in brain, which approached significance (p=0.0588, t test, parametric data distribution). Viral titers in the lungs were not significantly affected by pixatimod treatment (**Fig 4B**). In view of the robust nature of this model, with rapid effects such as massive weight loss and high level of lethality, these data indicate a remarkable prophylactic anti-SARS-CoV-2 effect of pixatimod.

## Discussion

The current COVID-19 pandemic illustrates the critical need to develop both effective vaccines and therapeutics for emerging viruses; established antiviral agents appear to have limited utility against SARS-CoV-2. Owing to their use as a means of cell attachment by many viruses, HS represents an ideal broad-spectrum antiviral target (*2*). Binding of a viral protein to cell-surface HS is often the first step in a cascade of interactions that is required for viral entry and the initiation of infection (*29*). As HS and heparin contain the same saccharide building blocks and HS-binding proteins also interact with heparin, this drug is gaining attention beyond its anticoagulant properties in COVID-19 treatment (*29*). Here we demonstrate a direct mechanism of action of pixatimod on attenuating S1-RBD binding to ACE2. These data are supported by recent studies on heparin using native mass spectrometry (*30*), and also reveal for the first time the ability of HS mimetics to inhibit S1-RBD binding to ACE2.

Heparin has been shown to inhibit binding of SARS-CoV-2 spike protein to a human cell line (*31*), and to inhibit entry into human cells of pseudovirus carrying the SARS-CoV-2 spike protein (*10,32*). However, the question of whether therapeutic doses of heparins are effective for COVID-19 patients as an antiviral treatment awaits the outcome of clinical trials; bleeding complications are possible (*33*), though non-anticoagulant heparin or HS preparations could be deployed that reduce cell binding and infectivity without a risk of causing bleeding (*9,10*). However, HS mimetics offer additional advantages in comparison to heparin beyond simply reducing anticoagulant activity (9), most notably their ready availability at scale via synthetic chemistry production that addresses the well-known fragility of the heparin supply chain (*11*). As a clinical-stage HS mimetic, pixatimod provides better control over structure, molecular weight diversity (a single molecular entity), sulfation, purity and stability. Herein, we reveal a direct interaction of the clinical candidate pixatimod with the S1 spike protein RBD, supported by molecular modelling data. Pixatimod also inhibited the interaction of S1 RBD with Vero cells which express the ACE2 receptor. Moreover, infectivity assays, of two types (plaque reduction and cytopathic effect, Table 1) confirm pixatimod is a potent inhibitor of SARS-CoV-2 infection of Vero cells, at concentrations ranging from 0.8 to 13.8 µg/mL which are well within its known therapeutic range. Interestingly, we noted that the lipophilic steroid side chain of pixatimod was critical for its potency and is predicted from modelling to interact with S1-RBD. This unique feature, making it an unusual amphiphilic HS mimetic, has also been shown to confer virucidal activity against Herpes Simplex virus by disruption of the viral lipid envelope (*15*).

Pixatimod has only mild anti-coagulant activity, and has been administered i.v. to over 80 cancer patients, being well tolerated with predictable pharmacokinetics (PK) and no reports of heparin-induced thrombocytopenia (*12*). Further, cytotoxicity *in vitro* is low; we observed a CC_50_ >100 µM (>236 µg/mL) in Vero cells, consistent with cytotoxicity data on human cells (*34*). Importantly, the maximum plasma concentration (Cmax) of pixatimod following a single treatment of 100 mg in cancer patients is 29.5 µg/mL with a Cmin of 2.7 µg/mL measured one week following treatment (*12*), indicating that an equivalent dosing regimen should be sufficient to achieve antiviral activity in human subjects. The low anticoagulant activity of pixatimod is an advantage since it could be used as a direct antiviral agent in combination therapies with heparins, which are being used to treat coagulopathies observed in COVID-19 patients (*35*). It was also encouraging that pixatimod inhibition of multiple clinical isolates of SARS-CoV-2 was noted, demonstrating potential for widespread effectiveness. The presence of multiple binding sites for pixatimod in the Spike protein would suggest robustness against mutations that may arise later in pandemic and/or in the following coronavirus outbreaks. While recent widespread isolates with D614G spike mutants appear to be 2-3 fold more sensitive to the antiviral activity of pixatimod, caution needs to be taken in interpreting the data of the cytopathicity assay used to determine this activity as G614 isolates (at least QLD935) exhibited lower cytopathicity than D614 isolates.

Since SARS-CoV-2 is predominantly a respiratory infection, our data from human bronchial cells are directly supportive of high potency of pixatimod against infection of clinically relevant human cells. Further, the potential efficacy in humans is supported by our initial data from the established K18 hACE2 transgenic mouse model which demonstrated the ability to rescue the dramatic weight loss phenotype observed. Given the half-life of pixatimod in mice is 38h (*34*), compared to 141h in human (*13*), future studies should adopt repeat-dose schedules to identify the optimal schedule in models of SARS-CoV-2 infection. We also suggest that direct intranasal delivery of pixatimod would be worth investigation, in view of recent positive data on the delivery of nebulized heparin for treatment of ARDS (*36*) which has significance for potential application of heparin in COVID-19 treatment (21).

Why Pixatimod should have such a dramatic effect in nasal turbinates and brain, but not lungs in the K18 hACE2 model is unclear but may reflect very high lung (supra-physiological) K18-driven hACE2 receptor expression in the lungs in this model (*27*) and/or pharmacokinetics (*34*). Nevertheless the results are encouraging, as the initial infection in humans occurs in the upper respiratory tract, with ARDS developing only after infection has over time spread down into the lungs (*37*). This progression is not recapitulated in the mouse model where fulminant lung infection requires direct inoculation of virus into the lungs via the intranasal route. Infection of the brain may occur via the olfactory epithelium (*38*) so the reduced infection of brain may reflect the reduced infection of nasal turbinates. Although CNS involvement in COVID-19 in humans is now well documented^32^, the fulminant brain infection seen in this mouse model does not seem to be a feature of human disease.

It is notable that there are multiple potential mechanisms of action of pixatimod against SARS-CoV-2 (summarised in **Fig 5**), including direct inhibition of HS-Spike-ACE2 interactions via complex and multiple interactions with amino acids in RBD. Notably, the latter contrasts with the more restricted epitopes typical of antibodies which are observed to be subject to mutational escape potentially requiring vaccine redesign or use of multivalent strategies (*39*). In this respect we speculate that heparin mimetics like pixatimod may prove important therapeutic tools for initial responses to escape variants. Beyond this. pixatimod also has immunomodulatory effects which may alleviate some of the immunopathologies associated with moderate-severe COVID-19 disease. Pixatimod also inhibits the pro-inflammatory enzyme heparanase (*34*) and has been demonstrated to suppress IL-6 in inflammatory (pancreatitis) and viral (Ross River virus) animal models (*14, 18*). Moreover, it blocks the heparanase-dependent invasion of macrophages into tumours in mouse cancer models (*35*) which may be relevant to invasion of monocytes and macrophages into the lungs associated with severe COVID-19 disease (*40*). Vaccinia virus has recently been shown to rely on host heparanase to degrade HS in order to spread to distant sites (*41*), revealing a role for heparanase in the progression of disease that may also apply for SARS-CoV-2 in COVID-19. Notably, increased plasma heparanase activity is associated with COVID-19 (*42*). Thus multiple additional beneficial effects of pixatimod might be anticipated from its heparanase inhibitory properties.

**Figure 5:**
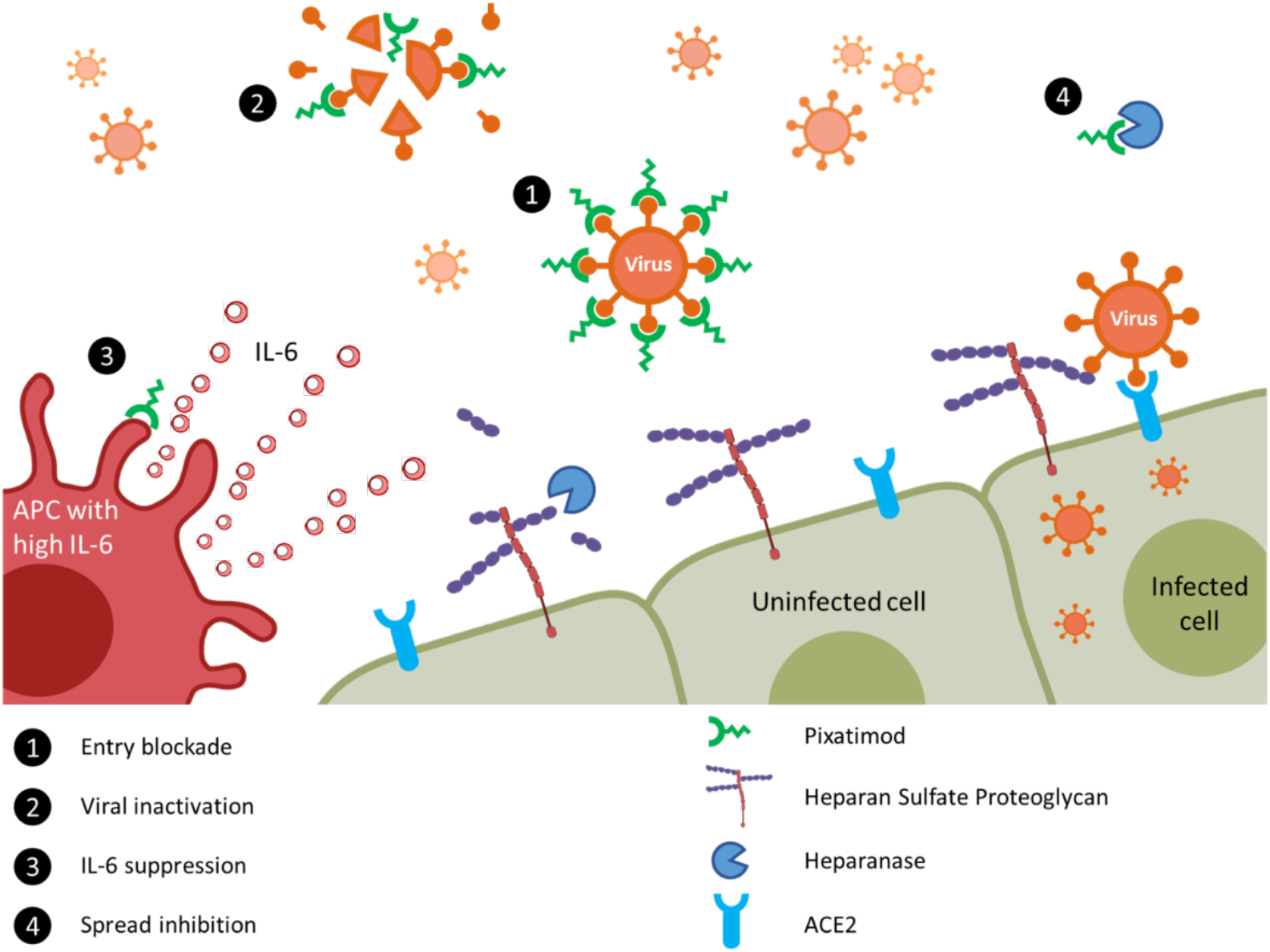
Proposed multi-modal mechanisms of pixatimod activity against SARS-CoV-2 and other viruses. The principal mode of action demonstrated here is that pixatimod acts as a decoy receptor^1^, blocking S1-RBD binding to HS co-receptors and inhibiting viral attachment to host cells, thus blocking viral infection. Additional potential modes of action include: [2] virucidal activity of pixatimod, dependent upon the cholestanol moiety^15^, which may lead to degradation and permanent inactivation of SARS-CoV-2 virus particles; [3] suppression of IL-6 secretion by antigen presenting cells, primarily macrophages^14^; and [4] blocking viral escape from host cells by inhibiting heparanase which otherwise promotes viral escape by cleaving HS receptors^41,42^.

Based on the data presented here, pixatimod has potent antiviral activity against SARS-CoV-2 at therapeutically relevant concentrations both *in vitro* and *in vivo*, and against relevant human bronchial cells. These activities are in addition to the known heparanase-inhibitory and immunomodulatory properties of pixatimod which may further support the host response to COVID-19 infection. Collectively this provides a strong rationale to justify entry of pixatimod into clinical trials for COVID-19. Furthermore, we have demonstrated the first proof-of-concept for employing HS mimetics against SARS-CoV-2 with implications for wider future applications of this class of broad-spectrum antivirals, potentially against SARS-CoV-2 escape variants, other HS-binding viruses and also those that may emerge as future global threats.

## Materials and Methods

### Computational methods

The crystal structure of the RBD-ACE2 complex (PDB ID: 6LZG) was retrieved from the RCSB Protein Data Bank. Structures were stripped of water molecules, ACE2 chain and any cofactors and/or ligands present. UCSF Chimera was used to edit the structure and for visualisation. Without prior knowledge of the pixatimod binding site, one molecule of the ligand was placed in the simulation system containing the protein, solvent and ions and molecular dynamics (MD) simulations were performed for 600 ns using the AMBER16 package. Such unguided simulations, as reviewed before (*43*), have been used to predict the binding sites on a protein’s surface and drive the design of new ligands. All the MD simulations were carried out using the pmemd.cuda module (*44*) of the AMBER 16 molecular dynamics package and the analyses were performed using the cpptraj module of AmberTools16 (*45*). Simulation systems were set up by placing the spike RBD domain at the centre of the octahedral simulation box (with an extension of at least 12 Å from each side). Pixatimod was randomly placed in the box. This was followed by addition of TIP3P water (*46*) and Na^+^ ions for neutralising the charge of the system. Proteins were parameterized using the Amber99SB-ildn force field (*47*) whereas Glycam-06 (version j) (*48*) and Lipid14 (*49*) force fields were used for the sulfated tetrasaccharide and cholestanol moieties of pixatimod, respectively. Four replicates of the unguided simulations were performed (4 × 600 ns). Periodic boundary conditions were applied, and the time step was set to 2 fs. The electrostatic energy was calculated with the particle mesh Ewald (PME) method. SHAKE constraints were applied on the bonds involving hydrogen. A cut-off of 12 Å was applied to the Lennard-Jones and direct space electrostatic interactions with a uniform density approximation included to correct for the long-range van der Waals interactions.

The system was first minimized without electrostatics for 500 steps, then with a restraint of 25 kcal/(mol Å^2^) applied on the protein and pixatimod. This minimization was followed by 100-ps MD simulation with 25 kcal/(mol Å^2^) positional restraints applied on the protein and ligand, and the temperature was slowly increased from 0 to 300 K. Then, followed by 500 steps of steepest descent cycles followed by 500 steps of conjugate gradient minimization, and 50-ps equilibrations with a restraint force constant of 5 kcal/(mol Å^2^) in the protein and ligand, followed by final 2 ns equilibration without restraints to equilibrate the density. The first few steps were all carried out at constant volume followed by at least 600 ns production MD simulation at 300 K (Langevin dynamics, collision frequency: 5/ps) with 1-atm constant pressure. Trajectories were collected and data analyses such as RMSD, RMSF and free energy of binding were performed on the last 30000 frames. The binding free energy and pairwise residue contributions (*50*) were calculated using the Molecular mechanics-Generalized Born (GB) equation (MM/GBSA) procedure implemented in AmberTools16. The details of this method have been extensively reviewed (*51*). The polar solvation energy contribution was calculated by using GB^OBC II^ (igb= 5) (*52*). The value of the implicit solvent dielectric constant and the solute dielectric constant for GB calculations was set to 80 and 1, respectively. The solvent probe radius was set to 1.4 Å as default. The entropy calculation is computationally expensive and therefore not performed for the purposes of this study.

### Expression of His-tagged recombinant SARS-CoV-2 S1 RBD in E. coli

Residues 330−583 of the SARS-CoV-2 spike protein (GenBank: MN908947) were cloned upstream of a N-terminal 6XHisTag in the pRSETA expression vector and transformed into SHuffle® T7 Express Competent *E. coli* (NEB, UK). Protein expression was carried out in MagicMediaTM *E. coli* Expression Media (Invitrogen, UK) at 30°C for 24 hrs, 250 rpm. The bacterial pellet was suspended in 5 mL lysis buffer (BugBuster Protein Extraction Reagent, Merck Millipore, UK; containing DNAse) and incubated at room temperature for 30 mins. Protein was purified from inclusion bodies using IMAC chromatography under denaturing conditions. On-column protein refolding was performed by applying a gradient with decreasing concentrations of the denaturing agent (from 8M Urea). After extensive washing, protein was eluted using 20 mM NaH_2_PO_4_, pH 8.0, 300 mM NaCl, 500 mM imidazole. Fractions were pooled and buffer-exchanged to phosphate-buffered saline (PBS; 140 mM NaCl, 5 mM NaH_2_PO_4_, 5 mM Na_2_HPO_4_, pH 7.4; Lonza, UK) using Sephadex G-25 media (GE Healthcare, UK). Recombinant protein (termed EcS1-RBD) was stored at -4°C until required.

### Expression of S1-RBD in mammalian cells

Secreted RBD-SD1 (termed mS1-RBD) was transiently produced in suspension HEK293-6E cells. A plasmid encoding RBD-SD1, residues 319−591 of SARS-CoV-2 S were cloned upstream of a C-terminal HRV3C protease cleavage site, a monomeric Fc tag and an His_8x_ Tag were a gift from Jason S. McLellan, University of Texas at Austin. Briefly, 100 mL of HEK293-6E cells were seeded at a cell density of 0.5 × 106 cells/ml 24hr before transfection with polyethyleneimine (PEI). For transfection, 100 μg of the ACE2 plasmid and 300 μg of PEI (1:3 ratio) were incubated for 15 min at room temperature. Transfected cells were cultured for 48 hr and fed with 100 mL fresh media for additional 48 hr before harvest. RBD-SD1was purified by HiTrap Protein G HP column (GE Healthcare, US) pre-equilibrated in PBS and eluted with 0.1 M glycine (pH 2.7). Purity of proteins was evaluated by Coomassie staining of SDS-PAGE gels, and proteins were quantified by BCA Protein Assay Kit (Thermo Scientific).

### Secondary structure determination of SARS-CoV-2 S1 RBD by circular dichroism spectroscopy

The circular dichroism (CD) spectrum of the SARS-CoV-2 S1 RBD in PBS was recorded using a J-1500 Jasco CD spectrometer (Jasco, UK), Spectral Manager II software (JASCO, UK) using a 0.2 mm path length, quartz cuvette (Hellma, USA). All spectra were obtained using a scanning of 100 nm/min, with 1 nm resolution throughout the range λ = 190 - 260 nm and are presented as the the mean of five independent scans, following instrument calibration with camphorsulfonic acid. SARS-CoV-2 S1 RBD was buffer-exchanged (prior to spectral analysis) using a 10 kDa Vivaspin centrifugal filter (Sartorius, Germany) at 12,000 g, thrice and CD spectra were collected using 21 μL of a 0.6 mg/mL solution in PBS, pH 7.4. Spectra of heparin (unfractionated porcine mucosal heparin, Celsus) were collected in the same buffer at approximately comparable concentrations, since this is a polydisperse material. Collected data were analysed with Spectral Manager II software prior to processing with GraphPad Prism 7, using second order polynomial smoothing through 21 neighbours. Secondary structural prediction was calculated using the BeStSel analysis server (*53*).

To ensure that the CD spectral change of SARS-CoV-2 S1 RBD in the presence of pixatimod did not arise from the addition of the compound alone, a difference spectrum was analysed. The theoretical CD spectrum that resulted from the arithmetic addition of the CD spectrum of the SARS-CoV-2 S1 RBD and that of pixatimod differed from the observed experimental CD spectrum of SARS-CoV-2 S1 RBD mixed with compound alone. This demonstrates that the change in the CD spectrum arose from a conformational change following binding to pixatimod (Supplementary Materials, Fig S3).

### Differential scanning fluorimetry

Differential scanning fluorimetry (DSF) was conducted on mammalian expressed mS1-RBD (1 μg) in PBS pH 7.6 and 1.25 X Sypro Orange (Invitrogen) to a total well volume of 40 μL in 96-well qPCR plates (AB Biosystems). Unfractionated porcine mucosal heparin (Celsus) or pixatimod (10 μg) were introduced to determine the effect on the thermal stability of, mS1-RBD using an AB biosystems StepOne plus qPCR machine, employing the TAMRA filter setting. Melt curve experiments were performed following a 2-minute initial incubation at 25 °C, with succeeding 0.5 °C increments every 30 s up to a final temperature of 90°C. Control wells containing H_2_O, heparin or pixatimod (10 μg) without mS1-RBD (1 μg) also employed to ensure a change in the melt curve was solely a result of protein-ligand interactions and interactions with Sypro Orange. Smoothed first derivative plots (9 neighbours, 2^nd^-order polynomial, Savitxky-Golay) were constructed using Prism 8 (GraphPad). T_m_ values were calculated using MatLab softaware (R20018a, MathWorks) and ΔT_m_ values determined from the difference between the T_m_ of RBD alone or in the presence of heparin or pixatimod.

### Cell binding of S1 RBD

African green monkey Vero kidney epithelial cells (Vero E6) were purchased from ATCC. Cells were maintained at 50-75% confluence in DMEM supplemented with 10% foetal bovine serum, 20 mM L-glutamine, 100 U/mL penicillin-G and 100 U/mL streptomycin sulfate (all purchased from Gibco/ThermoFisher, UK). Cells were maintained at 37 °C, in 5% CO_2_ and plated into 96-well cell culture plates at 1000 cells/well in 100 μL of maintenance medium. Cells were allowed to adhere overnight. Medium was aspirated and wells were washed 3x with 200 μL calcium, magnesium-free PBS (CMF-PBS, Lonza, UK). Cells were fixed with 100 μL 10% neutral buffered Formalin (Thermofisher, UK) for 10 minutes at room temperature, then washed 3x with 200 μL CMF-PBS. 100 μL CMF-PBS was added to each well and plates were stored at 4 °C until use. Before use, wells were blocked with 200 μL CMF-PBS + 1% BSA (Sigma-Roche, UK) for 1 hour at room temperature, and washed 3x with 200 μL CMF-PBS + 0.1% Tween-20 (PBST, Sigma-Roche, UK) followed by 2x with 200 μL CMF-PBS. His-tagged S1-RBD (50 μg/mL) and compounds at indicated concentrations were added to each well in 25 μL PBST + 0.1% BSA as indicated. Wells were incubated for 1 hour at room temperature with rocking. Wells were washed 3x with 200 μL PBST and 2x with 200 μL CMF-PBS. Binding of His-tagged S1-RBD was detected with Alexa Fluor 488 anti-his tag antibody (clone J095G46, Biolegend, UK) 1:5000 in 25 μL PBST + 0.1% BSA per well. Wells were incubated in the dark for 1 hour at room temperature with rocking. Wells were washed 3x with 200 μL PBST and 2x with 200 μL CMF-PBS. Fluorescence was read at Ex. 485:Em 535 on a Tecan Infinite M200Pro plate reader. Results are presented as normalized mean (where 0 is the fluorescence without added S1-RBD, and 1 is the fluorescence with 50 μg/mL S1-RBD; ± %CV, n=3).

### Competition ELISA for S1 RBD binding to ACE2

High binding 96 well plates (Greiner) were coated with 3 μg/mL streptavidin (Fisher) in 50 mM sodium carbonate buffer pH 9.6 (50 μL/ well) for 1 hour at 37 °C. Plates were washed 3 times with 300 μL PBS, 0.2% Brij35 (PBSB) and blocked with 300 μL PBSB + 1% casein for 1 hour at 37 °C. Plates were washed a further 3 times with 300 μL PBSB prior to the addition of 100 ng/mL BiotinylatedACE2 (Sino Biological) in PBSB + 1% casein (50 μL/ well) and incubated for 1 hour at 37 °C. Plates were again washed 3 times with 300 μL PBSB prior to the addition of 50 μL/well mS1-RBD (5μg/mL) in PBSB + 1% casein, which had been pre-incubated for 30 minutes at room temperature with or without varying concentrations of heparin or pixatimod (100-0.7 μg/mL) in separate tubes. Plates were incubated for 1 hour at 37 °C to allow for mS1-RBD-ACE2 binding and were subsequently washed with 300 μL/well PBSB. Bound mS1-RBD was detected by incubation with 0.5 μg/mL Rabbit-SARS-CoV-2 (2019-nCoV) Spike RBD Antibody (Stratech) in PBSB + 1% casein (50 μL/well) for 1 hour at 37 °C. Following a further 3 washes with PBSB plates were incubated for 30 minutes at 37 °C with horseradish peroxidase conjugated Donkey anti-Rabbit IgG diluted 1:1000, v/v in PBSB + 1% casein (Bioledgend). Plates were washed a final 5 times with 300 μL PBSB before being developed for 10 minutes with 3,3’,5,5’-tetramethylbenzidine prepared according to the manufacturer’s instructions (Fisher). Reactions were stopped by the addition of 20 μL 2M H_s_SO_4_ and plates were read at λ =450 nm using a Tecan Infinate M200 Pro mulit-well plate reader (Tecan Group). Control wells containing no biotinylated ACE2 were employed to ensure binding was specific.

### Live SARS-CoV-2 virus assays

SARS-CoV-2 Victoria isolate (GISAID accession, EPI_ISL_406844): a plaque reduction assay was performed with the SARS-CoV-2 Victoria/01/2020 (passage 3) isolate, generously provided by The Doherty Institute, Melbourne, Australia at P1, was diluted to a concentration of 1.4 × 10^3^ pfu/mL (70 pfu/50µL) in minimal essential media (MEM) (Life Technologies, California, USA) containing 1% (v/v) foetal calf serum (FCS) (Life Technologies) and 25 mM HEPES buffer (Sigma) and mixed 50:50 with pixatimod dilutions, in a 96-well V-bottomed plate. The plate was incubated at 37 °C in a humidified box for 1 hour to allow the virus to be exposed to pixatimod. The virus-compound mixture was transferred onto the wells of a washed 24-well plate that had been seeded with Vero E6 cells [ECACC 85020206] the previous day at 1.5 × 10^5^ cells/well. The virus-compound mixture was left to adsorb for an hour at 37°C, then plaque assay overlay media was applied (MEM containing 1.5% carboxymethylcellulose (Sigma, Dorset, UK), 4% (v/v) FCS and 25 mM HEPES buffer). After incubation at 37 °C in a humidified box, for 5 days, plates were fixed overnight with 20% (v/v) formalin/PBS, washed with tap water and then stained with methyl crystal violet solution (0.2% v/v) (Sigma) and plaques were counted. Compound dilutions were performed in either duplicate or quadruplicate. Compound dilutions and cells only were run in duplicate, to determine if there was any cell cytotoxicity. A mid-point probit analysis (written in R programming language for statistical computing and graphics) was used to determine the amount (μg/mL) required to reduce SARS-CoV-2 viral plaques by 50% (PRNT50) compared with the virus only control (n=5). An internal positive control for the PRNT assay was run in triplicate using a sample of heat-inactivated human MERS convalescent serum known to neutralise SARS-CoV-2 (National Institute for Biological Standards and Control, UK).

SARS-CoV-2 DE-Gbg20 isolate (GISAID accession under application): Plaque reduction assay for SARS-CoV-2 clinical isolate DE-Gbg20 from Sweden was performed in a similar manner, except for the virus and the pixatimod (fivefold decreasing concentrations at a range 100-0.16 µg/ml) were diluted in DMEM supplemented with 2% heat-inactivated FCS, and 100 U of penicillin and 60 µg/ml of streptomycin (DMEM-S). The virus (100 PFU) and pixatimod (fivefold decreasing concentrations at a range 100 – 0.16 µg/ml) were mixed and incubated for 30 min in humidified atmosphere comprising 5% CO_2_ (CO_2_ incubator). The mixtures were then transferred to Vero cells (ATCC CCL-81) and following incubation with cells for 90 min in the CO_2_ incubator, the methylcellulose overlay was added. Three separate experiments each with duplicates were performed.

A cytopathic effect assay was performed with the SARS-CoV-2 DE-Gbg20 isolate and Vero cells (ATCC) plated at 2 × 10^4^ per well in 96-well plates the day prior to the experiment. Serial fivefold dilutions of pixatimod in DMEM supplemented with 2% heat-inactivated FCS, and 100 U of penicillin and 60 µg/mL of streptomycin (DMEM-S) were incubated with 100 TCID50 of SARS-CoV-2 isolate DE for 20 min in humidified atmosphere comprising 5% CO_2_ (CO_2_ incubator). The final concentrations of pixatimod were in a range 0.075 µg/mL to 47.2 µg/mL. The cells were rinsed once with 50 µL of DMEM-S, and then 200 µL of the virus-pixatimod mixtures were added to each well with cells (in quadruplicates). After incubation of the virus-pixatimod mixtures with cells for 3 days in the CO_2_ incubator, the cells were inspected under a microscope for the presence of virus induced cytopathic effect where complete protection of cells were denoted as “+” while a partial protection (∼50% of cells showing no cytopathic effect) was recorded as “+/-”. The 50% end-point (EC_50_) was computed by the Reed and Muench method.

SARS-CoV-2 QLD02 (GISAID accession EPI_ISL_407896) and QLD935 (GISAID accession EPI_ISL_436097) clinical isolates from Australia: A cytopathic effect assay was carried out as described above for the DE-Gbg20 isolate, with 10 ffu/well and 3 days incubation. In this assay, Vero E6 cells were plated at 2 × 10^4^ per well in 96-well plates the day prior to experiment. Serial five-fold dilutions of pixatimod in DMEM supplemented with 2% heat-inactivated FCS, and 100 U of penicillin and 60 µg/mL of streptomycin (DMEM-S) were incubated with 10 foci forming units of SARS-CoV-2 QLD02 or QLD935 isolate and incubated for 30 min in humidified atmosphere comprising 5% CO_2_ (CO_2_ incubator). The cells were rinsed once with 50 µL of DMEM-S, and then 200 µL of the virus-pixatimod mixtures were added to each well with cells (in triplicates). After incubation of the virus-pixatimod mixtures with cells for 3 days in the CO_2_ incubator, the cells were fixed with 4% PFA and then stained with crystal violet. Then crystal violet was released by methanol and OD at 595nm was measured to quantify cell viability (protection from infection). The EC_50_ was then calculated using GraphPad Prism.

### Cytotoxicity assays

The assay was performed as described previously (*17*). Briefly, Vero cells (ATCC, 2 × 10^4^ cells/well) were seeded in 96 well cluster plates to become nearly confluent at the day of the experiment. The cell growth medium was then removed and 100 µL of serial fivefold dilutions of pixatimod in DMEM-S (ranging from 0.09 to 236 µg/mL) were added to cells. Following incubation of cells with pixatimod for 3 days in the CO_2_ incubator, 20 µL of the MTS salt containing CellTiter 96 Aqueous One Solution reagent (Promega, Madison, WI) was added and incubated for further 1-2 h at 37 °C. The absorbance was recorded at 490 nm against a background of 650 nm. Two separate experiments each in duplicates were performed and the results are expressed as percentage of absorbance value detected with pixatimod relative to control cells.

### Human bronchial cell infection assays

The hTERT transformed bronchial epithelial cell line BCi-NS1.1 (*23,24*) expanded in PneumaCult-Ex Plus Basal Medium supplemented with Pneumacult Ex Plus supplements, hydrocortisone, nystatin and penicillin–streptomycin. BCi-NS1.1 cells were grown at an Air Liquid Interface (ALI) in PneumaCult-ALI Basal Medium supplemented with Pneumacult ALI supplement hydrocortisone, PneumaCult ALI maintenance supplement, heparin, nystatin and penicillin–streptomycin (all from Stemcell Technologies) at 37°C in 5% CO_2_. BCi-NS1.1 cell ciliation was observed by microscopy and cells were differentiated and ciliated after 4 to 5 weeks at ALI. Transepithelial electrical resistance (TER) was monitored weekly using an EVOM Voltohmmeter (World Precision Instruments) and cells with a TER ≥1000 WΩm2 were used.

SARS-CoV-2 strain BetaCoV/ Australia/VIC01/2020 was used for infection: 50,000 plaque-forming units (MOI=0.1-0.2, obtained from Public Health England and propagated in Vero E6 cells for no more than two passages before use) were preincubated with either 4, 40 and 400 μg/ml of Wockhardt heparin or 0.4, 4 and 40 μug/ml of pixatimod for 1 hr at 37 ° C in a total volume of 200 μl (or an equivalent volume of saline was added apically in the untreated controls). BCi-NS1.1 cells (4 to 8 weeks after ALI) after being washed three times with HBSS were apically infected with the preincubated virus and compounds. After 2h the virus-compound solution was removed from the apical side and washed twice each time with 200ul HBSS. The last wash was collected for plaque assays and 50 μl stored in Qiazol. Similarly at 72 h a further 200 μl HBSS wash was performed and collected as at 2h. Cells were also harvested with QIAzol (QIAGEN) at 2 h after the HBSS wash and at 72 h post-infection for RNA extraction.

Plaque assay was performed to quantify extracellular virus released in the supernatant of infected ALI. Vero E6 cells were seeded at 2.5 × 10^5^ cells/well in a 12-well plate and left for a period of 24 hours in DMEM supplemented with 10% foetal bovine serum (FBS), glutamine and 50 U/ml penicillin-streptomycin at 37°C in 5% CO_2_. Cells were washed once with infection medium (serum-free DMEM supplemented with 25 mM HEPES) and 100 μl from washes of infected ALI cultures added to wells in 10 fold serial dilutions. After a 1 hour incubation at 37°C in 5% CO2, infectious supernatants were removed and a 1.5ml overlay of 1 x DMEM supplemented with 4% FBS, 25 mM HEPES and 0.6% (w/v) cellulose (Sigma; cat no 435244) was added. Plates were incubated at 37°C and 5% CO2 for 72 hours before removing the overlay, fixing with 8% formaldehyde in PBS, and staining with 0.1% (w/v) crystal violet in a solution of 20% (v/v) ethanol.

RNA was isolated from cell lysates using standard phenol–chloroform extraction, and reverse transcribed to cDNA using a High Capacity cDNA Reverse Transcription kit (Thermo Fisher Scientific) following the manufacturer’s instructions. Taqman gene expression assays for N2 and N3 regions of the SARS-CoV-2 N gene were made (ThermoFisher) following CDC specifications (Division of Viral Diseases, National Center for Immunization and Respiratory Diseases, Centers for Disease Control and Prevention, Atlanta, GA, USA, 20 January 2020 copy).

To measure viral shedding in the apical side, gene expression was quantified using the 2^−ΔCt^ method. For determination of relative changes of RNA within the cells the average Ct values of both SARS-CoV-2 N2 and N3 gene RNA were normalised against the house keeping gene B2M and shown as fold changes compared to the untreated cells using the 2^−ΔΔCt^ method.

### K18-hACE2 mouse experiments

Mouse experiments are approved by the QIMR Berghofer MRI Biosafety Committee and Animal Ethics Committee (project P3600), and are conducted in accordance with the “Australian Code for the care and use of animals for scientific purposes” as defined by the National Health and Medical Research Council of Australia. Work was conducted in a dedicated suite in a biosafety level-3 (PC3) facility at the QIMR Berghofer MRI (Australian Department of Agriculture, Water and the Environment certification Q2326 and Office of the Gene Technology Regulator certification 3445). Heterozygous K18 hACE2-transgenic mice (The Jackson Laboratory, Bar Harbour, ME, USA) were bred in-house by crossing with C57BL/6J mice (Animal Resources Center, Canning Vale, WA, Australia). DNA from tail tips was isolated using Extract-N-Amp Tissue PCR kit (Sigma) and PCR genotyping undertaken as described (The Jackson Laboratory. Genotyping protocols database. B6.Cg-Tg(K18-ACE2)2Prlmn/J. Stock No: 034860. Protocol 38276), except using primers Forward – 5’-CTTGGTGATATGTGGGGTAGA-3’, reverse 5’ CGCTTCATCTCCCACCACTT-3’ (recommended by NIOBIOHN, Osaka, Japan).

Female 6-8 week old K18 hACE2 mice (n=4 per group) were treated once with 16 mg/kg of pixatimod in 200 µl saline i.p. on the day before infection. Control K18 hACE2 mice received 200 µl saline on the day before infection. Age and gender matched C57BL/6J mice (n=4) were included as a drug toxicity control and received 16 mg/kg of pixatimod in 200 µl saline i.p., but were not infected with virus.

hCoV-19/Australia/QLD02/2020 (QLD02) (GISAID Accession ID; EPI_ISL_407896) virus was used to inoculate K18 hACE2 mice (n=4 per group) intranasally with 10^4^ CCID_50_ of virus in 50 μl medium while under light anaesthesia; 3% isoflurane (Piramal Enterprises Ltd., Andhra Pradesh, India) delivered using The Stinger, Rodent Anesthesia System (Advanced Anaesthesia Specialists/Darvall, Gladesville, NSW, Australia). Mice were weighed daily and euthanized on day 5 post-infection using CO_2_. Tissues were fixed for histology or were weighed and then homogenised using four ceramic beads at 6000 rpm twice for 15 s (Precellys 24 Homogeniser, Bertin Instruments, Montigny-le-Bretonneux, France). After centrifugation for 30 mins, 2000 g at 4°C, virus titres in supernatants were determined by CCID_50_ assays using Vero E6 cells.

### Statistical analysis

Experimental data are presented as means ± SD, SEM or CV as noted. Statistical analyses were performed using analysis of a two-tailed Student’s *t* test with GraphPad Prism (GraphPad Software) unless otherwise noted. Differences were considered statistically significant if the *P* value was less than 0.05. Statistical analysis for the K18-hACE2 mouse work was performed using IBM SPSS Statistics for Windows, Version 19.0 (IBM Corp., Armonk, NY, USA). The t-test was used when the data was deemed parametric, i.e. difference in variances was <4, skewness was >2 and kurtosis was <2. Otherwise the non-parametric Kolmogorov-Smirnov test was used.

## Acknowledgments

The authors would like to thank Zucero Therapeutics for provision of pixatimod (PG545) and Queensland Health Forensic & Scientific Services, Queensland Department of Health for provision of QLD02 and QLD935 SARS-CoV-2 isolates.

## Author contributions

J.E.T., V.F., M.A.S. and K.D. conceived the project. S.E.G., C.J.M-W., N.S.G., J.A.T., T.T.L., C.M.S., M.V.H., K.R.B., N.C., M.J.E., K.N., J.S., Y.X.S., A.A.A., N.M., J.D.J.S., M.C., D.W., P.R.Y., A.A.K., M.A.L., E.A.Y., R.K., R.L.M., Y-H.C., Z.Y., E.T.,M.A.S., T.M.A.W. and A.S. designed and conducted the experiments and undertook analyses. V.F., E.H., K.D. M.W.C., J.S., T.B., M.A.S. and J.E.T. analyzed results and prepared the manuscript.

## Data and materials availability

All data needed to evaluate the conclusions in the paper are present in the paper and/or the Supplementary Materials. Additional data related to this paper may be requested from the authors.

## Supplementary Materials

### The clinical-stage heparan sulfate mimetic pixatimod (PG545) potently inhibits SARS-CoV-2 virus via disruption of the Spike-ACE2 interaction

Scott E. Guimond, Courtney J. Mycroft-West, Neha S. Gandhi, Julia A. Tree, Karen R. Buttigieg, Naomi Coombes, Kristina Nyström, Joanna Said, Yin Xiang Setoh, Alberto A. Amarilla, Naphak Modhiran, De Jun Julian Sng, Mohit Chhabra, Daniel Watterson, Paul R. Young, Alexander A. Khromykh, Marcelo A. Lima, Edwin A.Yates, Richard Karlsson, Yen-Hsi Chen, Yang Zhang, Edward Hammond, Keith Dredge, Miles W. Carroll, Edward Trybala, Tomas Bergström, Vito Ferro, Mark A. Skidmore and Jeremy E. Turnbull

**Table S1:** Amino acids at position 614 in Spike protein of SARS-CoV-2 isolates.

| Isolate | Amino acids at position 614 in Spike |
| --- | --- |
| VIC01 | D |
| QLD02 | D |
| QLD935 | G |
| DE-Gbg20 | G |

**Figure S1.**
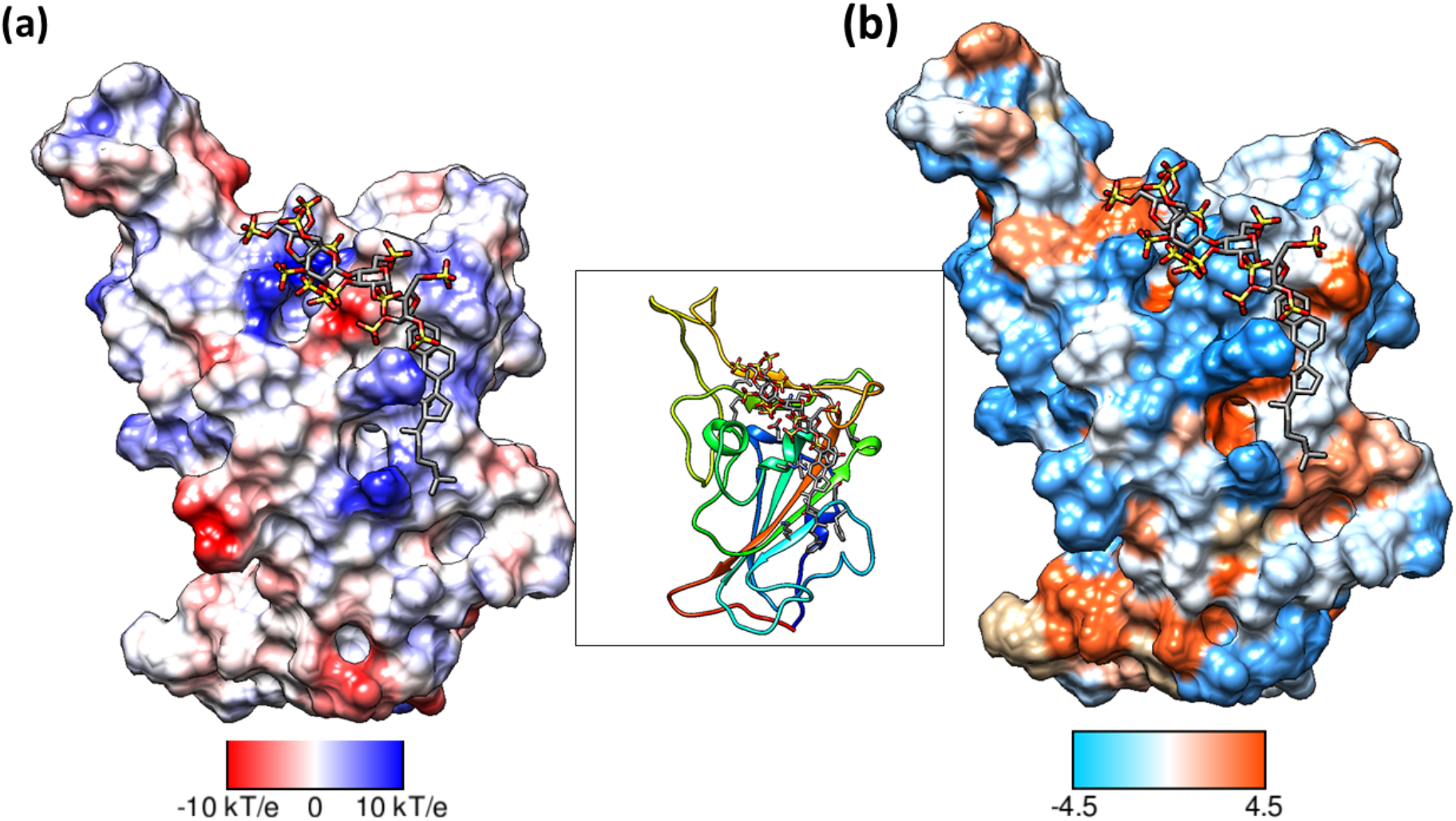
An alternate binding mode of pixatimod on the S1 RBD presented an unfavourable total binding free energy. Surfaces are oriented in the same direction as shown in the ribbon diagram in the inset. (a) Coulombic Surface Coloring defaults: ε = 4r, thresholds ± 10 kcal/mol·e were used. Blue indicates surface with basic region whereas red indicates negatively charged surface. (b) The hydrophobic surface was coloured using the Kyte-Doolittle scale wherein blue, white and orange red colour indicates most hydrophilic, neutral and hydrophobic region, respectively. UCSF Chimera was used for creating surfaces and rendering the images. Hydrogens are not shown for clarity.

**Figure S2.**
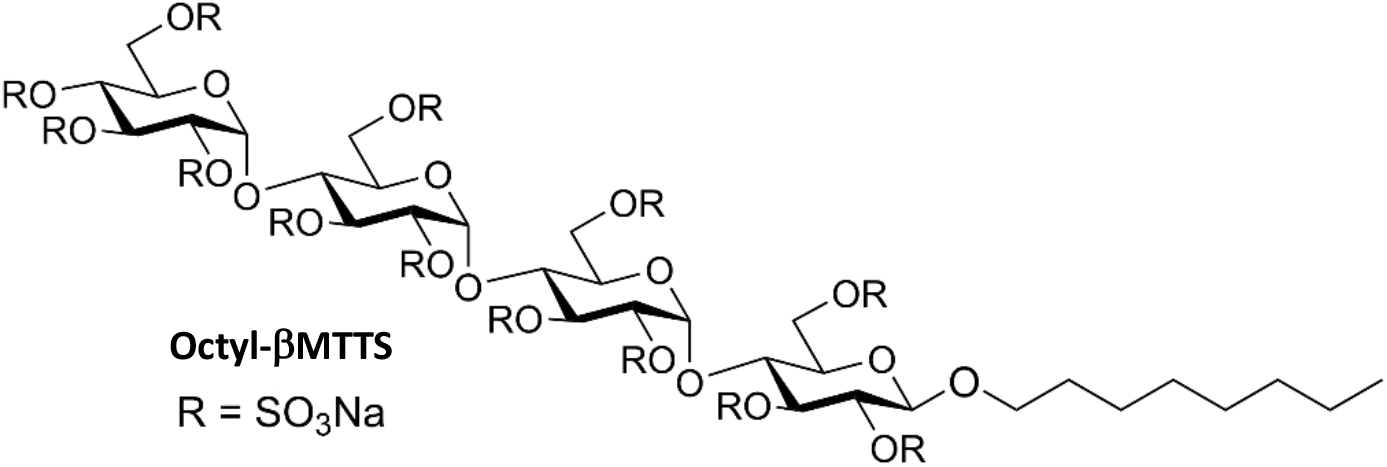
Structure of octyl β-maltotetraoside tridecasulfate, an analogue of pixatimod without the steroid side chain.

**Figure S3:**
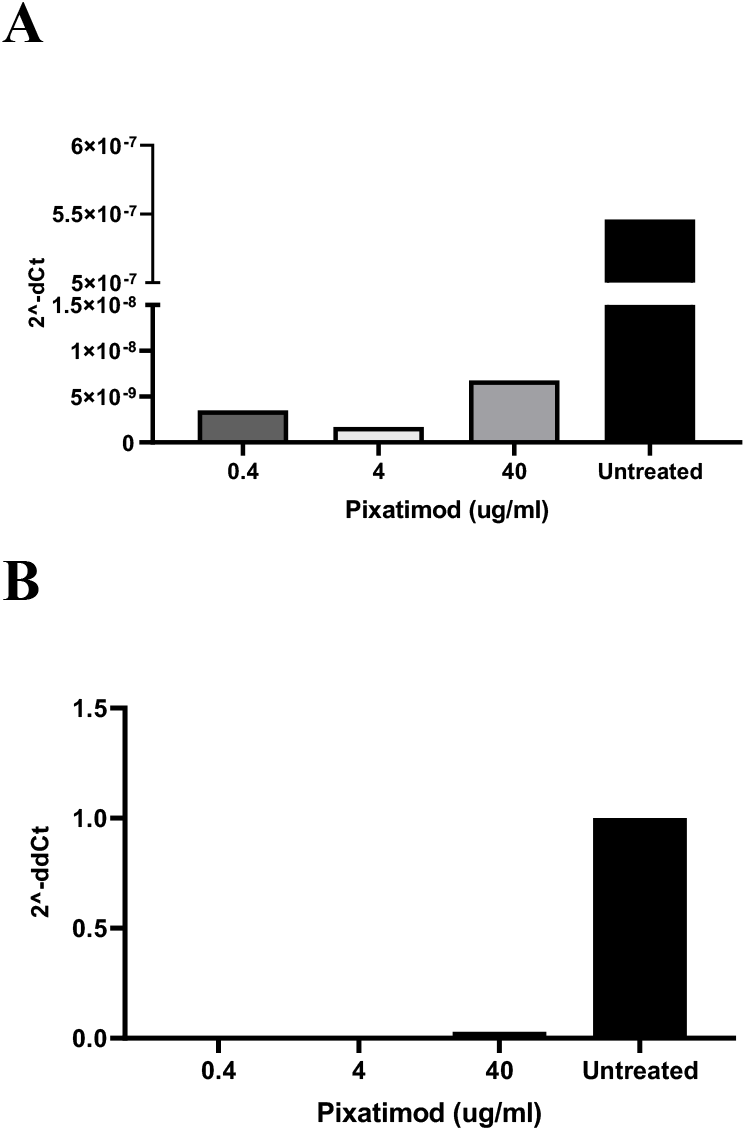
Pixatimod inhibits infection of human bronchial epithelial cells with SARS-CoV-2 virus. Live virus infectivity assays of bronchial airway epithelial cells BCi-NS1.1, grown in an air liquid interface (ALI), were performed as described in Methods for the SARS-CoV-2 isolate VIC01. Viral shedding in the apical side (panel A) and viral load in infected cells (panel B) was measured by RT-PCR of viral RNA, as described in Methods. (representative data shown).

**Figure S4:**
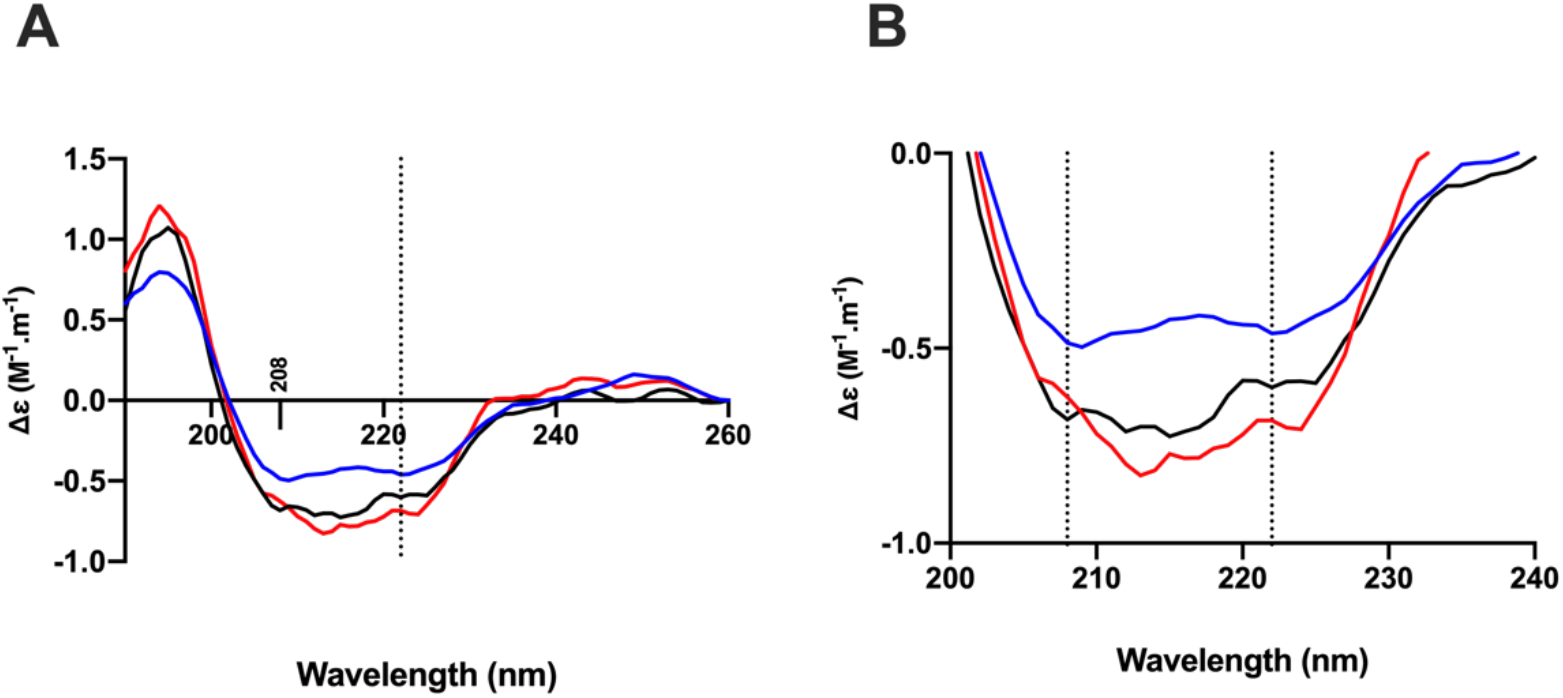
The conformational change of the SARS-CoV-2 S1 RBD observed in the presence of pixatimod by circular dichroism (CD) spectroscopy. (A). Circular dichroism spectra (190 - 260 nm) of SARS-CoV-2 S1 RBD alone (black solid line) and pixatimod (blue solid line) in PBS, pH 7.4. The red line represents the sum of the two individual spectra. Vertical dotted line indicates 222 nm (B) Details of the same spectra expanded between 200 and 240 nm. Vertical dotted lines indicate 222 nm and 208 nm.

